# Development of a genome-scale metabolic model of *Clostridium thermocellum* and its applications for integration of multi-omics datasets and strain design

**DOI:** 10.1101/2020.04.02.022376

**Authors:** Sergio Garcia, R. Adam Thompson, Richard J. Giannone, Satyakam Dash, Costas D. Maranas, Cong T. Trinh

**Author notes:** Corresponding author: 1512 Middle Dr, DO432, Department of Chemical and Biomolecular Engineering, University of Tennessee, Knoxville, TN 37996, USA.

## Abstract

Solving environmental and social challenges such as climate change requires a shift from our current non-renewable manufacturing model to a sustainable bioeconomy. To lower carbon emissions in the production of fuels and chemicals, plant biomass feedstocks can replace petroleum using microorganisms as catalysts. The anaerobic thermophile *Clostridium thermocellum* is a promising bacterium for bioconversion due to its capability to efficiently degrade untreated lignocellulosic biomass. However, the complex metabolism of *C. thermocellum* is not fully understood, hindering metabolic engineering to achieve high titers, rates, and yields of targeted molecules. In this study, we developed an updated genome-scale metabolic model of *C. thermocellum* that accounts for recent metabolic findings, has improved prediction accuracy, and is standard-conformant to ensure easy reproducibility. We illustrated two applications of the developed model. We first formulated a multi-omics integration protocol and used it to understand redox metabolism and potential bottlenecks in biofuel (e.g., ethanol) production in *C. thermocellum*. Second, we used the metabolic model to design modular cells for efficient production of alcohols and esters with broad applications as flavors, fragrances, solvents, and fuels. The proposed designs not only feature intuitive push-and-pull metabolic engineering strategies, but also novel manipulations around important central metabolic branch-points. We anticipate the developed genome-scale metabolic model will provide a useful tool for system analysis of *C. thermocellum* metabolism to fundamentally understand its physiology and guide metabolic engineering strategies to rapidly generate modular production strains for effective biosynthesis of biofuels and biochemicals from lignocellulosic biomass.

## 1 Introduction

Global oil reserves will be soon depleted,^1^ and climate change could become a major driver of civil conflict.^2^ These challenges to security and the environment need to be addressed by replacing our current non-renewable production of energy and materials for a renewable and carbon neutral approach.^3^ The gram-positive, thermophilic, cellulolytic, strict anaerobe *C. thermocellum* is capable of efficient degradation of lignocellulosic biomass to produce biofuels and biomaterial precursors, making this organism an ideal candidate for consolidated bioprocessing (CBP), where saccharification and fermentation take place in a single step.^4^ However, its complex and poorly understood metabolism remains the main roadblock to achieve industrially competitive titers, rates, and yields of biofuels such as ethanol^5^ and isobutanol.^6^

For the past decade, significant efforts have been dedicated to characterize and manipulate *C. thermocellum*’s central metabolism, due to increasing interest in developing this organism as a CBP manufacturing platform for biofuels production.^7^ *C. thermocellum* possesses atypical central metabolism, characterized by the important roles of pyrophosphate and ferredoxin,^8^ which makes redirection of both carbon and electron flows for biofuel production challenging to achieve. Specifically, *C. thermocellum* has various reactions to regulate intracellular concentration levels of NADH, NADPH, and reduced ferredoxin. These cofactors are used as electron donors with high specificity throughout metabolism. To maintain redox balance, *C. thermocellum* also possesses several hydrogenases to oxidize these reduced cofactors to molecular hydrogen that is secreted by the cell. Removal of these hydrogenases through deletion of *ech* (encoding the ferredoxin-dependent hydrogenase, ECH) and *hydG* (associated with the bifurcating hydrogenase, BIF, and bidirectional hydrogenase, H2ASE syn) was successfully applied to increase ethanol yield by electron rerouting.^9^ Thompson *et al.*^10^ characterized the Δ*hydG*Δ*ech* strain in depth by flux analysis of its core metabolism, concluding that the major driver for ethanol production was redox rather than carbon balancing. In particular, the conversion of reduced ferredoxin to NAD(P)H is likely the most rate limiting step. In a subsequent study, Lo *et al.*^11^ over-expressed *rnf* (encoding the ferredoxin-NAD oxidoreductase, RNF) in the Δ*hydG*Δ*ech* strain that is expected to enhance NADH supply, but did not achieve improved ethanol yield.

In an attempt to redirect carbon and electron flows for enhanced ethanol production, Deng *et al.*^12^ manipulated the pyruvate node and malate shunt of *C. thermocellum*. By converting phosphoenolpyruvate (*pep*) to oxaloacetate (*oaa*) and then to pyruvate (*pyr*), this shunt can interchange one mole of NADPH with one mol of NADH generated from glycolysis. Interestingly, the authors noted that replacement of the malate shunt by alternative pathways not linked to NADPH increased ethanol production and carbon recovery but reduced amino acid formation, confirming the role of the malate shunt as an NADPH source in *C. thermocellum*.

Sulfur metabolism also plays a key role in redox metabolism of *C. thermocellum* and has been investigated for its role in ethanol production. Sulfate, a component of *C. thermocellum* media, serves as an electron acceptor, which is capable of oxidizing sulfate to sulfite and then sulfide. Thompson *et al.*^10^ demonstrated that the strain Δ*hydG*Δ*ech*Δ*pfl*, which cannot grow in conventional medium due to its inability to secrete hydrogen or formate, recovered growth by sulfate supplementation to the culture medium. More recently, Biswas *et al.*^13^ reported an increase in final sulfide concentration and over-expression of the associated sulfate uptake and reduction pathway in the Δ*hydG* strain, but did not observe a significant difference in final sulfide concentration in Δ*hydG*Δ*ech*. Remarkably, neither of the strains consumed cysteine from the medium, unlike the wild-type. Sulfide can be converted to cysteine by CYSS (Cysteine synthase) or homocysteine and then methionine by SHSL2 and METS (succinyl-homoserine succinate-lyase and methionine synthase), but the connection between the cessation of cysteine uptake and sulfate metabolism remains unclear.

Overall, the complex interactions of *C. thermocellum* metabolic pathways remain challenging to understand and engineer with conventional methods, and hence require a quantitative systems biology approach to decipher. To this end, several genome-scale metabolic models (GSMs) of *C. thermocellum* have been developed. The first GSM, named iSR432, was constructed for the strain ATCC27405 and applied to identify gene deletion strategies for high ethanol yield.^14^ This model was then further curated into iCth446.^15^ More recently, Thompson *et al.* developed the iAT601 genome-scale model^16^ for the strain DSM1313, which is genetically tractable.^17^ The iAT601 model was used to identify genetic manipulations for high ethanol, isobutanol, and hydrogen production,^16^ and to understand growth cessation prior to substrate depletion observed under high-substrate loading fermentations that simulate industrial conditions.^18^ In addition to these core and genome-scale steady-state metabolic models, a kinetic model of central metabolism, k-ctherm118, was recently developed and used to elucidate the mechanisms of nitrogen limitation and ethanol stress.^15^ Due to the biotechnological relevance of the *Clostridium* genus, GSMs have also been developed for other species,^19^ including *C. acetubutylicum*,^20–26^ *C. beijerinckii*,^27^ *C. butyricum*,^28^ *C. cellulolyticum*,^22^ and *C. ljungdahlii*.^29^

In this study, we developed an updated genome-scale metabolic model of *C. thermocellum*, named iCBI655, with more comprehensive and precise metabolic coverage, enhanced prediction accuracy, and extensive documentation. This model is a human-curated database that coherently represents all the available genetic, genomic, and metabolic knowledge of *C. thermocellum* from both experimental literature and bioinformatic predictions. Furthermore, the model can be applied not only to enable metabolic flux simulation but also to provide a framework to contextualize disparate datasets at the system level. As a demonstration for the model application, we first developed a quantitative multi-omics integration protocol and used it to to fundamentally study redox metabolism and potential redox bottlenecks critical for production of biofuels (e.g., ethanol) in *C. thermocellum*. Furthermore, we used the model, in combination with the previously developed ModCell tool,^30,31^ to design modular (chassis) cells^32^ for alcohol and ester production.

## 2 Results

### 2.1 Development of an upgraded *C. thermocellum* genome-scale model named iCBI655

The iCBI655 model was developed using the published iAT601 model^16^ as a starting point. The model improvements include updated metabolic pathways, new annotation, and new extensive documentation. A detailed account of these changes can be found in the Supplementary Material 1. Here, we highlight the most relevant modifications.

#### 2.1.1 Modeling updates

To facilitate model usage and reduce human error, the identifiers of reactions and metabolites were converted from KEGG into BiGG human-readable form.^33^ Additionally, reaction and metabolite identifiers were linked to the modelSEED database^34^ that enables analysis through the KBase web interface.^35^ The gene identifiers and functional descriptions were updated to the most current annotation (NCBI Reference Sequence: NC 017304.1). Metabolite formulas and charges from the modelSEED database^34^ were included in the model and reactions were systematically corrected for charge and mass balance by the addition of protons and water.

#### 2.1.2 Metabolic updates

The automated construction process used in the previous model introduced several inconsistencies that were corrected in the current model. We removed reactions that were blocked and non-gene-associated, apparently introduced during automated gap-filling. Two notable examples are (i) the blocked selenate pathway which lacks experimental evidence (e.g., selenoproteins have not been found in *C. thermocellum*), and (ii) blocked reactions involving molecular oxygen (e.g., oxidation of Fe^2+^ to Fe^3+^) that are not possible in strict anaerobes like *C. thermocellum*. Furthermore, tRNA cycling reactions were unblocked by including tRNA into the biomass reaction.^36^ Metabolite isomers were examined and consolidated under the same metabolite identifier when possible, leading to the removal of duplicated reactions and the elimination of gaps. Transport and exchange reactions were updated to reflect the export of amino acids and uptake of pyruvate as observed during fermentation experiments.^37^

In terms of specific reactions, oxaloaceate decarboxylase was eliminated from the model in accordance with recent findings.^38^ The stoichiometries of pentose-phospate reactions, including se-doheptulose 1,7-bisphosphate D-glyceraldehyde-3-phosphate-lyase (FBA3) and sedoheptulose 1,7-bisphosphate ppi-dependent phosphofructokinase (PFK3 ppi), were corrected (according to experimental evidence^39^) from the previous model by ensuring mass balance and avoiding lumping multiple steps into one reaction. Transaldolase (TALA) was removed from the model due to lack of annotation for this gene in *C. thermocellum* and to also ensure PFK essentiality as observed experimentally (personal communication from Prof. Lynd’s lab at Dartmouth College).

Several modifications were also performed in key bioenergetic reactions. The reactions catalyzed by membrane-bound enzymes, including inorganic diphosphatase (PPA)^8^ and membrane-bound ferredoxin-dependent hydrogenase (ECH)^40^, were corrected to capture proton translocation. Furthermore, hydrogenase reactions were updated to ensure ferredoxin association for all cases and remove those reactions that do not involve ferredoxin and only use NAD(P)H as a cofactor based on our recent understanding of *C. thermocellum* metabolism.^13^ Gene-protein-reaction associations were updated to represent experimental knowledge. For instance, the hydrogenases BIF (CLO11313 RS09060-09070) and H2ASE (CLO1313 RS12830, CLO1313 RS02840) require the maturase Hyd (CLO1313 RS07925, CLO1313 RS11095, CLO1313 RS12830) to be functional, and the maturase itself requires all of its subunits to operate, which enables accurate representations of *hydG* deletion genotypes.^9^

Two hypothetical reaction modifications were introduced to ensure consistency with reported phenotypes. First, to enable growth without the need for succinate secretion, as observed in experimental data (Supplementary Material 2), the reaction homoserine-O-trans-acetylase (HSERTA) was added to enable methionine biosynthesis (essential for growth) without succinate formation. Although this reaction is not currently known to be associated with any gene in *C. thermocellum*, it is known to be present in other *Clostridium* GSMs.^29^ Next, the reaction deoxyribose-phosphate aldolase (DRPA) was removed to ensure correct lethality prediction of the Δ*hydG*Δ*ech*Δ*pfl* mutant strain as well as the correct prediction of growth recovery in this mutant by addition of external electron sinks such as sulfate or ketoisovalerate (Table 1). The correct prediction of Δ*hydG*Δ*ech*Δ*pfl*-associated phenotypes is critical to successfully use the model for computational strain design.^30,32,41–45^

**Table 1:** Comparison of mutant growth rates predicted by iAT601 and iCBI655. Experimental values are taken form Thompson *et al.*^10^; for some mutants whose growth recovery, not growth rate, was reported, they are presented with “+”.

| Gene deletions | Medium | Fraction of W.T. growth rate (%) |  |  |
| --- | --- | --- | --- | --- |
|  |  | *iAT601* | *iCBI655* | Experiment |
| *hydG* | MTC | 100 | 100 | 73 |
| *hydG-ech* | MTC | 85 | 85 | 67 |
| *hydG-pta-ack* | MTC | 100 | 100 | 48 |
| *hydG-ech-pfl* | MTC | 58 | 0 | 0 |
| *hydG-ech-pfl* | MTC + fumarate | 377 | 726 | 0 |
| *hydG-ech-pfl* | MTC + sulfate | 58 | 65 | + |
| *hydG-ech-pfl* | MTC + ketoisovalerate | 97 | 101 | + |

### 2.2 Comparison of iCBI655 against other genome-scale models

We compared iCBI655 with the previous GSMs of *C. thermocellum* and the highly-curated GSM iML1515 of the extensively studied bacterium *Escherichia coli* (Table 2). The increased number of genes in iCBI655 with respect to iAT601 cover a variety of functions, including hydrogenase chaperones, cellulosome and cellulase, ATP synthase, and transporters. Remarkably, iCBI655 has a smaller percentage of blocked reactions than iAT601, indicating higher biochemical consistency. The number of metabolites in iCBI655 is smaller than those in iAT601 mainly due to the removal of metabolites that did not appear in any reaction, duplicated metabolites (e.g., certain isomers), and blocked pathways added automatically during gap-filling without any gene association. *C. thermocellum* DSM1313 has 2911 protein coding genes, 22% of which is captured by iCB655, while *E. coli* MG1655 has 4240 genes, 35% of which is included in iML1515. Overall, iCBI655 increases the coverage of the metabolic functionality of *C. thermocellum* but remains far from the well studied *E. coli*.

**Table 2:** Comparison of all genome-scale metabolic models of *C. thermocellum* and the latest *E. coli* model.

|  | iSR432 | iCth446 | iAT601 | iCBI665 | iML1515 |
| --- | --- | --- | --- | --- | --- |
| Strain | ATCC27405 | ATCC27405 | DSM1313 | DSM1313 | MG1655 |
| Genes | 432 | 446 | 601 | 665 | 1515 |
| Metabolites | 583 | 599 | 903 | 795 | 1877 |
| Reactions | 632 | 660 | 872 | 854 | 2712 |
| Blocked reactions | 39.2% | 32.1% | 40.8% | 35.1% | 9.8% |
| Reference | [14] | [15] | [16] | This study | [46] |

### 2.3 Training of model parameters under diverse conditions

Growth and non-growth associated maintenance (GAM and NGAM) are parameters that capture the consumption of ATP towards cell division and homeostasis, respectively. These are known to be condition-specific; however, genome-scale models do not include a mechanistic description that allows to determine these ATP consumption rates as part of the simulation. Instead, GAM is incorporated into the biomass pseudo-reaction and NGAM has its own pseudo-reaction that hydrolyzes ATP at a rate tuned by constraint parameters.

To increase model prediction accuracy for various conditions, we trained GAM and NGAM parameters of iCBI655 using an extensive dataset of 28 extracellular fluxes (Supplementary Material 2) measured during the growth phase under different reactor configurations, carbon sources, and gene deletion mutants. This approach is based on the method used to train the iML1655 *E. coli* model.^46^ Remarkably, we observed highly linear trends under three different conditions, including chemostat reactor with cellobiose as a carbon source, chemostat reactor with cellulose as a carbon source, and batch reactor with either cellobiose or cellulose as carbon sources (Figure 1 a). This model training has led to improved growth rate prediction of iCBI655 as compared to iAT601 that has previously been trained with only a smaller dataset (Figure 1 b). Specifically, the iAT601 training dataset was limited to batch conditions; hence, the inaccurate predictions of iAT601 were observed for chemostat conditions (Figure 1 c).

**Figure 1:**
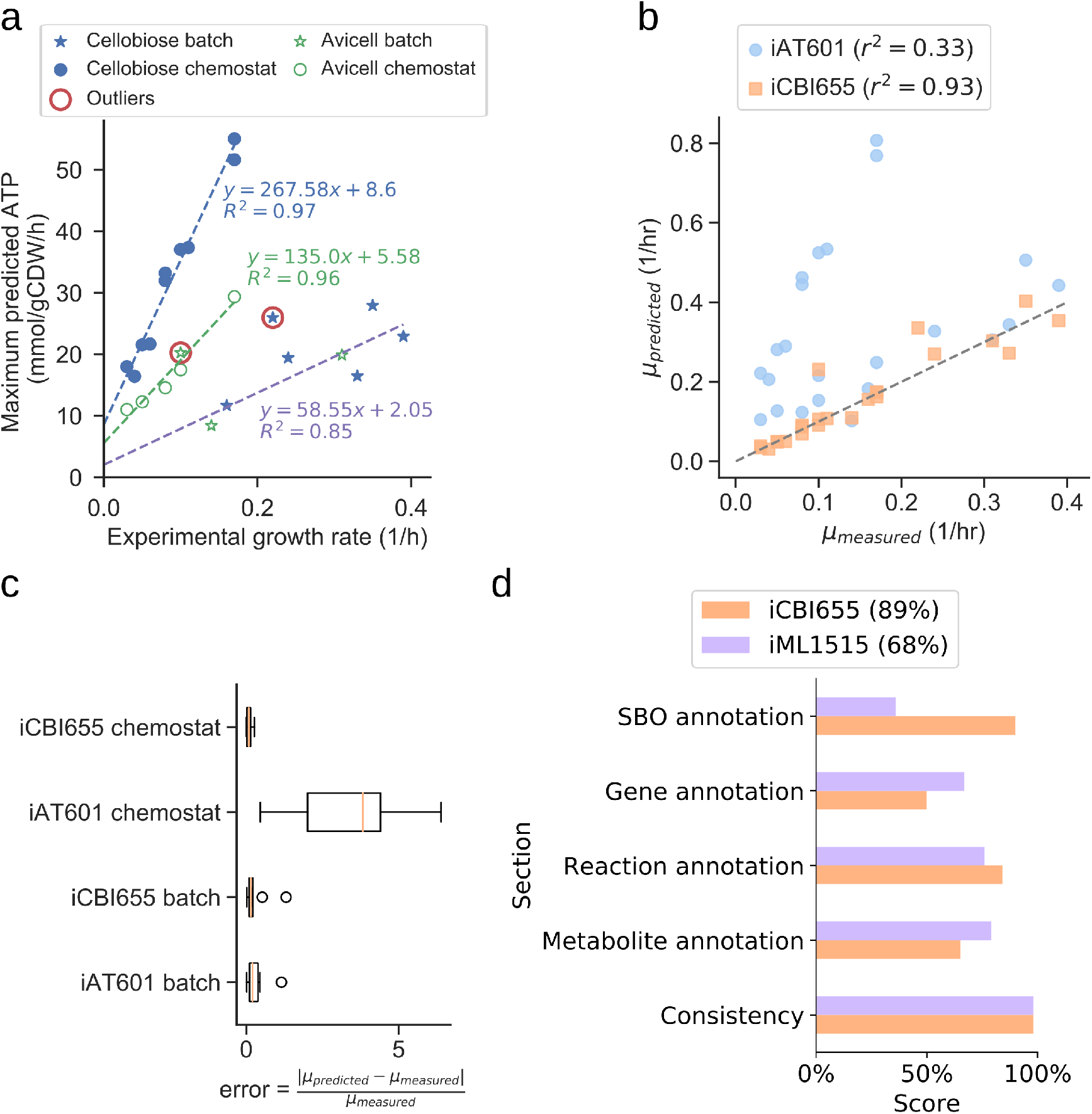
(**a**) Training of GAM and NGAM parameters. Discrete points correspond to experimental data. The slope of the linear regression function corresponds to the GAM, while the intercept corresponds to the NGAM. The data points circled as outliers were not included in any of the linear regression calculations. (**b**) Comparison of growth prediction error between iCBI655 and iAT601. Each maximum growth rate was predicted by constraining the models with the measured substrate uptake and product secretion fluxes (Supplementary Material 2). *r*^2^ corresponds to the Pearson correlation coefficient. (**c**) Error in growth predictions under batch and chemostat conditions. Predicted and measured growth rates correspond to the values included in **b**. (**d**) Scores provided by the quality control tool Memote^47^ for iCBI655 and iML1515. “Overall score” is shown in the legend.

### 2.4 Assessment of model quality and standard compliance with Memote

The field of metabolic network modeling suffers from a lack of standard enforcement and quality control metrics that limit model reproducibility and applicability. To address this issue, Lieven *et al.* recently developed the Memote framework that systematically tests for standards and best practices in GSMs.^47^ We applied Memote to both the iCBI655 and *E. coli* iML1515 models for comparison (Figure 1 d). This analysis produces five independent scores that assess model quality. The *consistency score* measures basic biochemical requirements, such as mass and charge balance of metabolic reactions, and it was near 100% for both models. Additionally, the different annotation scores quantify how many elements in the model contain relevant metadata. More specifically, the *systems biology ontology (SBO) annotation* indicates if an object in the model refers to a metabolite, reaction, or gene, while the respective *annotation scores* of these elements correspond to properties (e.g., name, chemical formula, etc.) and identifiers linking them to relevant databases (e.g., KEGG^48^ or modelSEED^34^). The *overall score* is computed as a weighted average of all the individual scores with additional emphasis on the *consistency score*. In summary, the high scores obtained by iCBI655 indicate the quality of the model and ensure its applicability for future studies.

### 2.5 Model-guided analysis of proteomics and flux datasets sheds light on redox metabolism critical for biofuel production in *C. thermocellum*

For the first application of the genome-scale metabolic model, we aimed to understand the complex redox metabolism and potential redox bottlenecks critical for enhanced biofuel production in *C. thermocellum*. We used the model as a scaffold to analyze proteomics and metabolic flux data collected for the *C. thermocellum* wild-type and Δ*hydG*Δ*ech* strains. The Δ*hydG*Δ*ech* mutant was engineered to redirect electron flow from hydrogen to ethanol by removal of primary hydrogenases.^9,10^ Previous studies of Δ*hydG*Δ*ech* based on analysis of secretion fluxes^10^ or omics^13^ suggested the presence of redox bottlenecks in this mutant but did not identify which specific path-ways and cofactors (i.e., NADH versus NADPH) are responsible. We aim to solve this problem through integrated and quantitative analysis of omics and fluxes at the genome scale.

#### 2.5.1 Development of fold change-based omics integration protocol

To perform the analysis, we formulated an omics integration protocol anchored in the quantification of fold change (FC) between case and control samples (Figure 2a). In this approach, we first compared FCs between simulated intracellular fluxes and measured omics data. Next, we identified *consistent reactions* with FCs of the same sign and different from zero in both measured proteomics and simulated fluxes for further analysis (Section 4.6).

**Figure 2:**
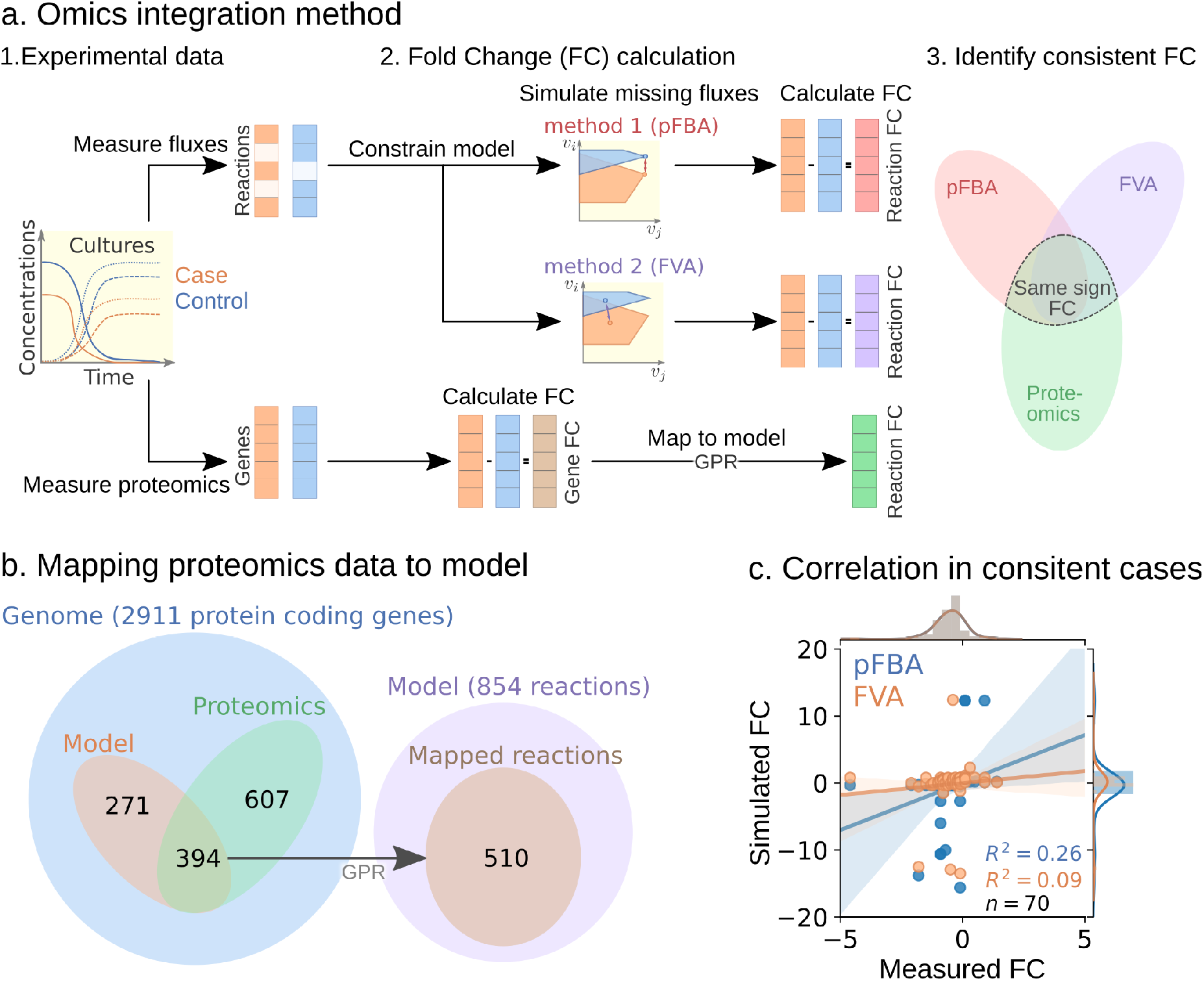
(**a**) Fold change-based multi-omics data integration and analysis protocol. (**b**) Mapping of proteomics data for the Δ*hydG*Δ*ech* case study to model reactions. (**c**) Correlation between measured and simulated fold changes (pFBA in blue and FVA in orange) for all 70 consistent reactions of the Δ*hydG*Δ *ech* case study.

To start the FC-based omics integration, we obtained measured FCs by mapping the measured proteomics data to 510 out of the 856 reactions in the model through the gene-protein-reaction (GPR) associations (Figure 2b). Then, we identified 70 consistent reactions by comparing measured FCs with two types of simulated FCs: i) parsimonious flux balance analysis (pFBA) that determines the most efficient flux distribution (assuming all enzymes are roughly as efficient) and ii) flux variability analysis (FVA) that identifies the feasible flux range of each reaction.

The Pearson correlation coefficients between simulated and measured FCs for the consistent reactions were 0.26 and 0.09 for pFBA and FVA, respectively (Figure 2c). In general, the FVA reaction flux ranges remained mostly unchanged, suggesting that pFBA is a better representation of actual metabolic fluxes as previously observed.^49^ The top consistent reactions with the highest proteomics FCs (Supplementary Material 3 - Table S1) belong primarily to the central metabolism of *C. thermocellum* (Figure 3). Interestingly, discrepancies in magnitude between flux and protein FCs for consistent reactions could be used to identify bottlenecks. For example, for a given enzyme a small increase in flux combined with a large increase in translation is an indicator of low catalytic efficiency. Similar comparisons between simulated flux and omics has previously been used to identify regulatory mechanisms.^50^ Overall, the identification and analysis of consistent reactions is an effective approach to gain certainty on the activity changes of metabolic pathways between conditions.

**Figure 3:**
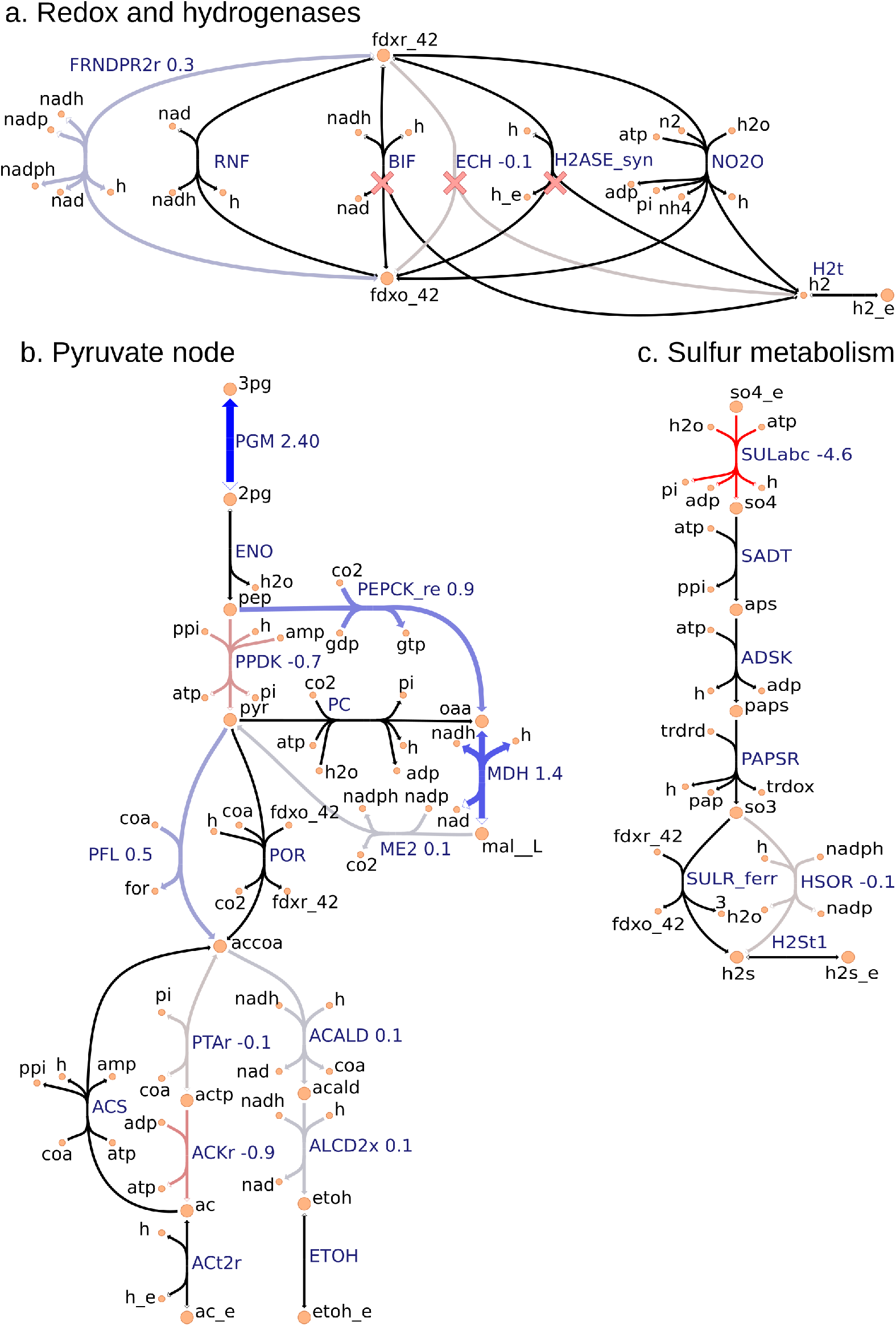
Metabolic map visualization using the iCBI Escher map. Values next to reaction labels correspond to proteomics fold change between Δ*hydG*Δ*ech* and wild-type strains only for the 70 consistent reactions identified by the omics analysis (Section 2.5). The labels of extracellular metabolites are appended with “_e”. Reactions marked with a red cross are deleted in Δ*hydG*Δ*ech*.

#### 2.5.2 FC-based omics integration reveals redirection of electron flow for NADPH supply inΔ*hydG*Δ*ech* strain

Our analysis reveals consistent indications of increased NADPH biosynthesis in the Δ*hydG*Δ*ech* mutant with respect to the wild-type across three major metabolic areas: i) an increased protein level of FRNDPR2r (also known as NFN) that converts one mol of reduced ferredoxin (*fdxr 42*) and one mole of NADH into two moles of NADPH (Figure 3a), ii) an increased protein level of all three malate shunt enzymes and a decreased protein level of the alternative route PPDK (Figure 3b); and iii) a decreased protein level of sulfur transporter and of HSOR that oxidizes sulfite into sulfide consuming NADPH (Figure 3c). These observations are consistent with the failure of *rnf* over-expression to enhance ethanol production,^11^ since RNF produces NADH but the key cofactor bottleneck seems to be NADPH. Furthermore, a direct look at the proteomics data revealed that RNF subunits (Clo1313 0061-Clo1313 0066) had a statistically significant decrease in protein levels of the mutant (Supplementary Material 2). The preference of Δ*hydG*Δ*ech* towards NADPH could be due to the cofactor specificity of the remaining redox balancing pathways (e.g., isobutanol), thermodynamics and protein cost constraints, or a combination of both. While the contribution of thermodynamic constraints is beyond the scope of this study, a recent analysis^51^ of the ethanol production pathway in *C. thermocellum* highlighted the importance of engineering strategies to increase NADPH, for instance, introduction of NADPH-linked GAPDH that converts glyceraldehyde-3-phosphate to 3-phospho-D-glyceroyl phosphate in glycolysis or overexpression of NADPH-FNOR that transfers electrons from reduced ferredoxin to NADPH. The contribution of alternative redox balancing pathway towards the increased NADPH biosynthesis will be examined next.

#### 2.5.3 Analysis of simulated fluxes reveals the role of NADPH in redox balancing

The analysis based on consistent reactions strongly indicates that NADPH production is important in the mutant to achieve redox balance. However, the pathways oxidizing NADPH remain unknown since not all reactions in the model could be mapped to proteomics measurements and carbon recovery was lower in the mutant strain.^10^ To identify these pathways, we examined the simulated fluxes of all reactions (instead of only consistent reactions) that differed in value between wild-type and mutant, and limited this search to exchange reactions and reactions that involve NADPH (Supplementary Material 3 - Table S2). These simulated fluxes predicted an increase in the isobutanol pathway, including keto-acid reductoisomerase (KARA1) that consumes NADPH and isobutanol secretion (EX ibutoh e). The isobutanol pathway can consume NADPH through several enzymes^6^ and has increased flux during overflow metabolism at high-substrate loading.^18,37^ The model also predicted a decrease in valine secretion (EX val L e), since the isobutanol pathway competes with the valine pathway after KARA1. Remarkably, this prediction is consistent with the lower valine secretion measured in Δ*hydG*Δ*ech*.^13^ A certain amount of NADPH is likely oxidized by the mutated alcohol-dehydrogenase enzyme observed after short adaptation in Δ*hydG* that is compatible with both NADH and NADPH.^9^ However, this feature is not captured by the model since in general gene knock-outs are simulated by blocking the associated reactions. Overall this analysis indicates that Δ*hydG*Δ*ech* likely increases isobutanol secretion to alleviate redox imbalance.

Taken altogether, model-guided data analysis illustrates the power of the model as contextualization tool and provides new insights into the redox bottlenecks present in *C. thermocellum* that are critical in the production of reduced molecules. The integration of omics and fluxes led to the resolution of NADPH as the key cofactor in redox imbalance of Δ*hydG*Δ*ech*, and it identified specific pathways that undergo major changes in protein levels, providing interesting target reactions for further engineering. Generally, the developed FC-based omics integration protocol can be applied to different omics data types due to its simplicity. The method does not require one to formulate or assume a quantitative relationship between omics measurements and simulated fluxes. Furthermore, fold change in biomolecule concentrations implemented in the method is currently much easier to measure in a quantitatively reliable manner for many molecules than case-specific absolute concentrations.

### 2.6 Model-guided design of modular production strains for biofuel synthesis

Another common application of genome-scale models is strain design.^30,32,41–45^ We used the iCBI655 model combined with the ModCell tool^31^ to design *C. thermocellum* modular production strains for efficient biosynthesis of alcohols and esters. Briefly, with ModCell we aim to design a modular (chassis) cell that can be rapidly combined with different pathway modules in a plug-and-play fashion to obtain modular production strains exhibiting target phenotypes with minimal strain optimization cycles. In this study, the target phenotype for modular production strains is growth-coupled to product synthesis (*wGCP*), that corresponds to the minimum product synthesis rate at the maximum growth rate. The design variables to attain the target phenotypes involve genetic manipulations of two types: i) reaction deletions, constrained by the parameter *α*, that corresponds to gene knock-outs; and ii) module reactions, constrained by the parameter *β*, that corresponds to reactions deleted in the modular cell but added back to specific modules to enhance the compatibility of a modular cell. Once these two parameters are specified, the solution to the problem is a set of Pareto optimal designs named Pareto front. In a Pareto optimal design, the performance (i.e., objective value) of a given module can only be increased at the expense of lowering the performance of another module. To characterize the practicality of each design, we say a modular cell is compatible with certain modules if the design objective is above a specific threshold (e.g., 0.5 in this study). Hence, the *compatibility* of a design corresponds to the number of compatible modules.

To design *C. thermocellum* modular cells, we first evaluated a range of design parameters *α* and *β* with an increasing number of genetic manipulations (Figure 4 a). As expected, increasing the number of deletions leads to more compatible designs, at the expense of more complexity in the implementation. We selected an intermediate point of *α* = 6, *β* = 0 for further analysis. This Pareto front is composed of 12 designs that can be clustered into two groups (Figure 4 b). The first group (e.g., designs 8, 3, and 9) are compatible with all products except butanol and its derived esters, whereas the second group (e.g., designs 2, 12, 1, and 10) have high objective values for butanol and its derived esters.

**Figure 4:**
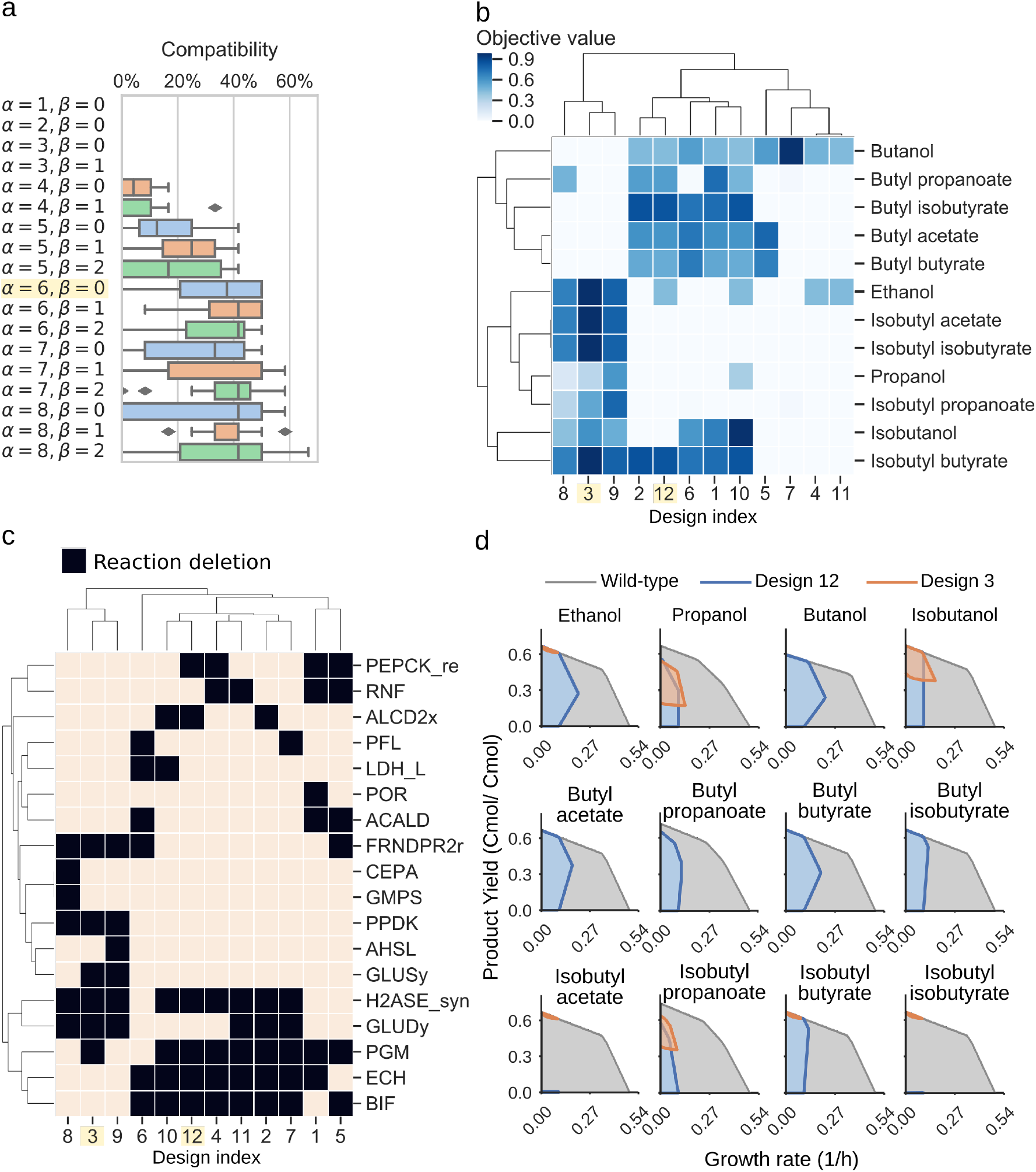
Proposed modular cell designs for biosynthesis of 12 alcohols and esters. (**a**) Module compatibility for various design parameters. (**b**) Pareto front for parameters *α* = 6, *β* = 0. Pareto set for parameters *α* = 6, *β* = 0. Reaction names and formulas are included in Supplementary Material 3 - Table S3. (**d**) Feasible phenotypic spaces for select designs.

To understand the characteristics of each group, we can inspect the deletions of each design (Figure 4 c). Designs 8, 3, and 9 all have in common H2ASE syn, GLUDy, PPDK, and FRNDPR2r deletion, while the last two deletions never appear in design 2, 12, 1, or 10. The majority of deletion targets are central metabolic reactions (Table 3). These include common targets such as deletion of hydrogenases that appear in the cluster of designs 10, 12, 4, 11, 2, and 7 with the Δ*hydG*Δ*ech* genotype discussed earlier or removal of reactions that form fermentative byproducts such as ALCD2x and ACALD (ethanol), PFL (formate), LDH L (lactate). Interestingly, ACKr or PTA (acetate) does not appear in this list, likely because acetate production can serve as a regulatory valve for redox metabolism, especially in a modular cell that must be compatible with products of diverse degrees of reduction.

**Table 3:** Reaction deletions sorted by appearance frequency (counts) in the designs of the Pareto front for *α* = 6, *β* = 0.

| ID | Name | Formula | Counts (%) |
| --- | --- | --- | --- |
| PGM | Phosphoglycerate mu-<br>tase | $2pg\_c \leftrightarrow 3pg\_c$ | 75 |
| H2ASE_syn | Bidirectional [NiFe] Hy-<br>drogenase (Fe-H2) | $h2\_c + nadp\_c \leftrightarrow h\_c + nadph\_c$ | 75 |
| ECH | (FeFe)-hydrogenase,<br>ferredoxin dependent,<br>membrane-bound | $2.0\ fdxr\_42\_c + 3.0\ h\_c \leftrightarrow 2.0\ fdxo\_42\_c + h2\_c + h\_e$ | 66.7 |
| BIF | Bifurcating Hydrogenase | $2.0\ fdxr\_42\_c + 3.0\ h\_c + nadh\_c \leftrightarrow 2.0\ fdxo\_42\_c + 2.0\ h2\_c + nad\_c$ | 66.7 |
| GLUDy | Glutamate dehydroge-<br>nase (NADP) | $glu\_L\_c + h2o\_c + nadp\_c \leftrightarrow akg\_c + h\_c + nadph\_c + nh4\_c$ | 50 |
| FRNDPR2r | Ferredoxin: nadp reduc-<br>tase (NFN) | $2.0\ fdxr\_42\_c + h\_c + nadh\_c + 2.0\ nadp\_c \leftrightarrow 2.0\ fdxo\_42\_c + nad\_c + 2.0\ nadph\_c$ | 41.7 |
| RNF | Ferredoxin:NAD oxi-<br>doreductase (membrane<br>bound) | $2.0\ fdxr\_42\_c + 2.0\ h\_c + nad\_c \leftrightarrow 2.0\ fdxo\_42\_c + h\_e + nadh\_c$ | 33.3 |
| PEPCK_re | Phosphoenolpyruvate<br>carboxykinase (GTP) | $co2\_c + gdp\_c + pep\_c \rightarrow gtp\_c + oaa\_c$ | 33.3 |
| ALCD2x | Alcohol dehydrogenase<br>(ethanol) | $acald\_c + h\_c + nadh\_c \rightarrow etoh\_c + nad\_c$ | 25 |
| ACALD | Acetaldehyde dehydroge-<br>nase (acetylating) | $accoa\_c + h\_c + nadh\_c \rightarrow acald\_c + coa\_c + nad\_c$ | 25 |
| PPDK | Pyruvate, phosphate dik-<br>inase | $amp\_c + 2.0\ h\_c + pep\_c + ppi\_c \rightarrow atp\_c + pi\_c + pyr\_c$ | 25 |
| GLUSy | Glutamate synthase<br>(NADPH) | $akg\_c + gln\_L\_c + h\_c + nadph\_c \rightarrow 2.0\ glu\_L\_c + nadp\_c$ | 16.7 |
| PFL | Pyruvate formate lyase | $coa\_c + pyr\_c \rightarrow accoa\_c + for\_c$ | 16.7 |
| LDH_L | L-lactate dehydrogenase | $h\_c + nadh\_c + pyr\_c \rightarrow lac\_L\_c + nad\_c$ | 16.7 |
| POR | Pyruvate-ferredoxin oxi-<br>doreductase | $coa\_c + 2.0\ fdxo\_42\_c + pyr\_c \rightarrow accoa\_c + co2\_c + 2.0\ fdxr\_42\_c + h\_c$ | 8.3 |
| CEPA | Cellobiose phosphorylase | $cellb\_c + pi\_c \rightarrow g1p\_c + glc\_D\_c$ | 8.3 |
| GMPS | GMP synthase | $atp\_c + nh4\_c + xmp\_c \rightarrow amp\_c + gmp\_c + 3.0\ h\_c + ppi\_c$ | 8.3 |
| AHSL | O-Acetyl-L-homoserine<br>succinate-lyase | $achms\_c + cys\_L\_c \leftrightarrow ac\_c + cyst\_L\_c + h\_c$ | 8.3 |

More interestingly, we also found important branch-point deletion reactions^52^ in central metabolism that have not yet been explored for strain design. Most prominently, these reactions include GLUDy, PEPCK re, and PPDK, which appear in 50%, 33%, and 25% of the designs, respectively (Table 3). Both PEPCK re and PPDK present two alternative routes that influence the ratio of NADPH to NADH, which is relevant to control metabolic fluxes though the specific dependencies of certain enzymes towards each redox cofactor. Since GLUDy consumes NADPH and is a key reaction in amino-acid metabolism, this enzyme and related ones (e.g., GLUSy) are interesting targets for practical implementation. We speculate the two product groups emerge likely because the butanol pathway strictly depends on NADH due to the reactions ACOAD1z (acyl-CoA dehydrogenase) and HACD1 (3-hydroxyacyl-CoA dehydrogenase), while the ethanol, propanol, and isobutanol path-ways are more flexible in their use of NADH or NADPH. The designs 8, 3, and 9 perform poorly with butanol, and are also the only ones containing PPDK deletion. This deletion forces pep to pyruvate flux through the malate shunt that converts NADH to NADPH. Engineering of the co-factor specificities of the butanol pathway could be used to build one modular cell compatible with all products under consideration.

Two representative designs from the groups mentioned earlier are 3 and 12. Their feasible growth and production phenotypes reveal a tight coupling between product formation and growth rate (Figure 4 d). This phenotype enables pathway optimization through adaptive laboratory evolution, as previously done for ethanol,^5^ overcoming one of the main challenges of *C. thermocellum* engineering that is optimization of enzyme expression levels. Hence, the proposed modular cells can also serve as platforms for pathway selection and optimization. In summary, this analysis demonstrates the potential of the model to identify non-intuitive metabolic engineering strategies that can be key to build effective modular platform strains for the production of biofuels and biochemicals in *C. thermocellum*.

## 3 Conclusions

In this study, we developed a genome-scale metabolic model of the biotechnologically relevant organism *C. thermocellum*. Model development followed standards and best practices to ensure reproducibility and accessibility. We demonstrated the enhanced predictions of the model for diverse fermentation conditions and gene lethality. Genome-scale models have a broad range of applications in systems biology, including metabolic engineering, physiological discovery, phenotype interpretation, and studies of evolutionary processes.^53,54^ To illustrate the model applications, we chose to tackle the challenge of disparate data integration and interpretation at the systems level. We developed a fold-change-based omics integration method for this purpose, and used it to identify routes in central metabolism that were selected to increase NADPH generation in the Δ*hydG*Δ*ech* strain. This analysis revealed the importance of NADPH cofactor over its alternatives and provided new engineering targets for enhanced biosynthesis of reduced products in *C. thermocellum*. We also illustrated the use of the model to design *C. thermocellum* modular cells, using the ModCell tool.^32^ The proposed designs cover C2 through C4 alcohols and their derived esters, which are key target molecules for renewable production with *C. thermocellum*.^55^ The proposed designs feature a combination of previously-explored and novel strategies to couple target metabolite production with cellular growth. Like the well-developed genome-scale models^46,56^ of the important organisms *Escherichia coli* and *Saccharomyces cerevisiae* broadly used for strain engineering both in academia^57^ and industry^58^, we anticipate the iCBI655 genome-scale model will also provide a versatile tool for systems metabolic engineering of *C. thermocellum*.

## 4 Methods

### 4.1 Model curation

The genome scale model iCBI655 was constructed from iAT601^16^ by following the standard GSM development protocol.^59^ Reaction and metabolite identifiers were mapped from KEGG to BiGG using the BiGG API.^33^ Metabolite charges were obtained from modelSEED when available, and otherwise calculated using the Chemaxon pKa plugin^60^ for a pH of 7.2.^59^ The biomass objective function was consolidated into one pseudo-reaction avoiding the use of intermediate pseudo-metabolites present in iAT601. Reactions were assigned with a confidence level based on a standard genome-scale model annotations.^59^

### 4.2 Metabolic flux simulations

Constraint-based metabolic network modeling^54^ is based on the feasible flux space, Ω_*k*_, defined by network stoichiometry and flux bounds that represent thermodynamic constraints and measured values:

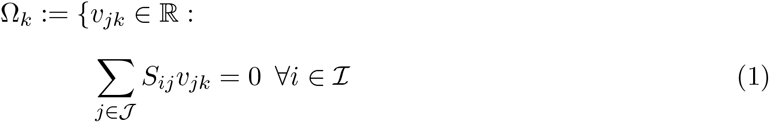

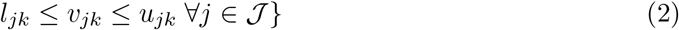

Here 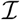 and 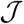 are the sets of metabolites and reactions in the model, respectively, and *v*_*j*_ is the metabolic flux (mmol/gCDW/h) through reaction *j*. Constraint (1) enforces mass balance for all metabolites in the network, where *S*_*ij*_ represents the stoichiometric coefficient of metabolite *i* in reaction *j*. Constraint (2) enforces lower and upper bounds *l*_*j*_ and *u*_*j*_, respectively, for each reaction *j* in the network.

In different simulation conditions, *k*, *S*_*ij*_ remains fixed given the structure of the network for all 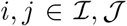. However, certain bounds *u*_*jk*_ and *l*_*jk*_ are modified to represent specific metabolic constraints. For example, to apply measured reaction fluxes such as in the case of GAM and NGAM calculation or the omics integration protocol (Section 4.6), *l*_*jk*_ and *u*_*jk*_ are specified using the experimentally measured average (*μ*_*jk*_) and standard deviation (*σ*_*jk*_), which for normally distributed samples with 3 replicates produces an interval with a confidence level above 90% (3-4). Similarly, to represent a certain gene deletion mutant *k*, the bounds are set to be *u*_*jk*_ = *l*_*jk*_ = 0 for the associated reaction *j*.

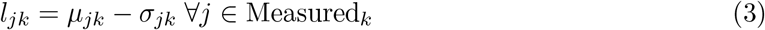

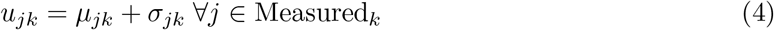

The feasible flux space Ω_*k*_ can be explored in different ways;^54,61^ for instance, an optimization objective is often defined to identify specific flux distributions 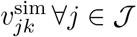:

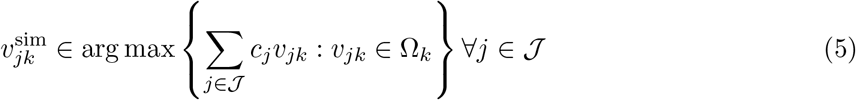

Here *c*_*j*_ is the coefficient of reaction *j* in the linear objective function, which is changed according to the simulation context. For example, to train GAM and NGAM (Figure 1a) the objective was set to maximize flux through the ATP hydrolysis reaction, i.e., *c*_*j*_ = 1 for *j* corresponding to ATP hydrolysis reaction, and 0 otherwise. To evaluate growth prediction accuracy (Figure 1b,c), the objective was set to maximize growth, i.e., *c*_*j*_ = 1 for *j* corresponding to growth pseudo-reaction and 0 otherwise.

### 4.3 Simulation of different growth environments

The model is configured to generally represent different medium and reactor conditions by modifying three features. The first feature involves model boundaries specifying which metabolites may enter the intracellular environment (i.e., present in the growth medium) or may exit the extra-cellular environment (i.e., secreted by *C. thermocellum*). This feature can be adjusted through *u*_*j*_ and *l*_*j*_ for exchange reactions. In our simulations, only essential metabolites required for *in silico* growth may be consumed and only commonly observed metabolites may be produced, unless otherwise noted. The second feature involves biomass objective function. iCBI655 contains 3 possible biomass reactions: i) BIOMASS CELLOBIOSE used for growth in cellobiose with cellulo-some constituting 2% of cell dry weight (CDW)^62^, ii) BIOMASS CELLULOSE used for growth on cellulose with cellulosome constituting 20% of CDW^62^, and iii) BIOMASS NO CELLULOSOME, a biomass function that does not consider cellulosome production and only used as a control. The combination of cellulosome and protein fractions accounts for 52.85% of the CDW in all cases.^10,14^ Cellobiose conditions were used in all simulations unless otherwise noted. The third feature involves GAM/NGAM. Three sets of these parameters are considered including batch, chemostat-cellulose, and chemostat-cellobiose, based on fitting the model to experimental data. Batch conditions were used in all simulations unless otherwise noted.

For growth on cellulose, the experimentally measured glucose equivalent uptake was represented in the model through the following pseudo-reactions: *3 glceq e* → *cell3 e*; *4 glceq e* → *cell4 e*; *5 glceq e* → *cell5 e*; *6 glceq e* → *cell6 e*. Here *cell3 e*, *cell4 e*, *cell5 e*, and *cell6 e* are cellodextrin polymers with 3, 4, 5, and 6 glucose monomers, respectively. These polymers can be imported inside the cell through the oligo-cellulose transport ABC system. The model is free to use any cellodextrin length, although higher lengths have higher ATP yield.^16,62^

### 4.4 Single-reaction deletion analysis to match experimentally observed phenotype

A core model of *C. thermocellum*^10^ correctly predicted the experimentally observed lethality of Δ*hydG*Δ*ech*Δ*pfl*; however, the iAT601 genome-scale model built by extension of this core model failed, suggesting that the genome-scale model has alternative active pathways leading to the false growth prediction *in silico*. To resolve this false positive prediction in iCBI655, we calculated the maximum growth rates for all possible additional single reaction deletions in the Δ*hydG*Δ*ech*Δ*pfl* mutant. This analysis resulted in three possible additional reaction deletions that are predicted to be lethal (i.e., maximum growth rate prediction below 20% of the simulated wild-type value^54^), including the removal of i) glycine secretion (EX gly e), ii) 5,10-methylenetetrahydrofolate oxidoreductase (MTHFC), and (iii) deoxyribose-phosphate aldolase (DRPA). For the first removal, addition of sulfate or ketoisovalerate in the growth medium of Δ*hydG*Δ*ech*Δ*pfl* fails to predict growth recovery as observed experimentally,^10^ making this option invalid. Likewise, the second removal is invalid because it makes PFL an essential reaction in the wild-type strain; however, experimental evidence demonstrates that Δ*pfl* mutant is able to grow.^63^ The last option was chosen since it correctly predicts growth recovery of Δ*hydG*Δ*ech*Δ*pfl* by sulfate or ketoisovalerate addition in the growth medium, and does not make PFL essential in the wild-type strain.

### 4.5 Model comparison

The *C. thermocellum* and *E. coli* models were obtained from their respective publications in SBML format. Blocked reactions are calculated by allowing all exchange reactions to have an unconstrained flux (i.e., *lb*_*j*_ = −1000, *ub*_*j*_ = 1000 ∀*j* ∈ *Exchange*). This enables the most general scenario which produces the smallest number of blocked reactions in each model. Additional details can be found in Supplementary Material 1.

### 4.6 Omics integration protocol

The omics integration protocol developed in this study consists of three steps: i) simulation of fold changes, ii) mapping of measured gene fold changes to reactions, and iii) comparison of measured and simulated fold changes.

#### 4.6.1 Calculation of simulated fold changes

To simulate metabolic fluxes, lower and upper bounds (2) are constrained according to experimental data as described in Section 4.2. Then, for the pFBA method, a quadratic optimization problem (6) is solved leading to a unique flux distribution 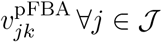

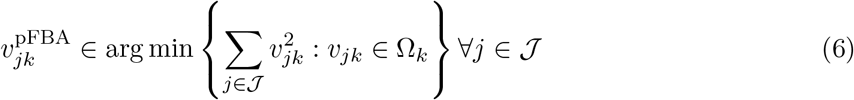

For the FVA method, a sequence of linear programming problems is solved where each flux is minimized (7) and maximized (8):

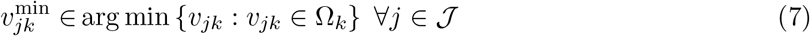

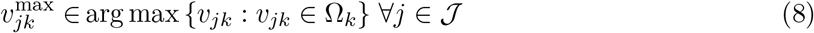

Note that for computation we applied the loop-less FVA method,^64,65^ as implemented in cobrapy,^66^ that introduces additional constraints in Ω_*k*_ to remove thermodynamically infeasible cycles from all feasible flux distributions.

FVA produces a flux range 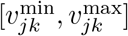 for each reaction 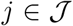. To compare between states *k* (e.g., wild-type and mutant), we define the *FVA center*, a scalar variable that indicates a change in this range (9).

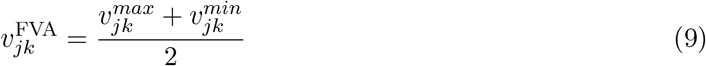

The FVA center, 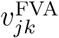, does not attempt to quantify the fraction of overlap between ranges nor to identify what type of shift might have occurred from all possible permutations, but simply provide an indicator of whether there is an upward shift (center increase) or downward (center decrease) between two conditions *k*. Unlike 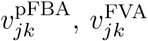 does not necessarily represent a feasible flux distribution of Ω_*k*_.

Finally, to determine the fold change for either pFBA or FVA simulated fluxes, the conventional procedure for fold change calculation in omics data is emulated. First, values are floored to avoid very large (or infinite) fold changes in cases with very small magnitude change. Given the minimum flux flooring value ∈ = 0.0001 a flooring function (10) is defined.

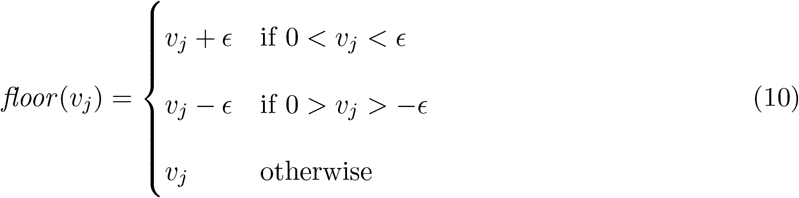

Then, the fluxes are normalized to the substrate uptake rate *v*_uptake, *k*_ and fold change is calculated in log_2_ space (11).

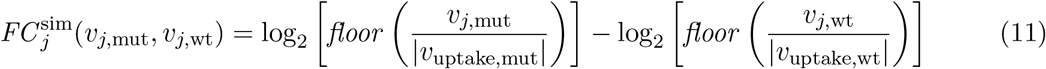

#### 4.6.2 Calculation of measured fold changes

Fold change between case and control samples, *FC*_*l*_, is calculated in log_2_ space for each gene 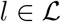, where 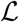 is the set of genes in the model. These gene fold changes can be mapped to metabolic reaction fold changes using the gene-protein reaction associations (GPR), given 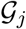 as the set of genes with *FC*_*l*_ ≠ 0 in the GPR of reaction *j*:

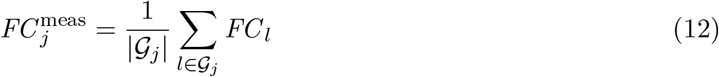

#### 4.6.3 Identification of consistent fold changes

A reaction *j* is said to have a consistent fold change if the measured fold change has the same sign of at least one of the simulated fold changes, more formally:

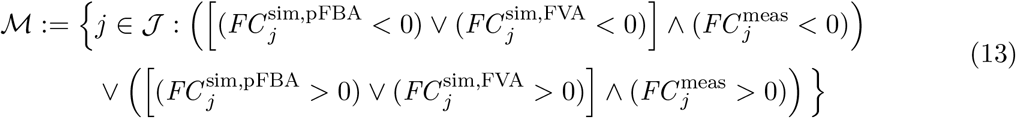

where 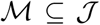 is the set of consistent reactions which is considered for further analysis and the simulated fold changes are re-defined for brevity (14-15).

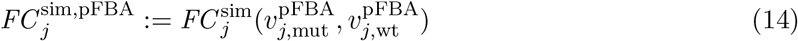

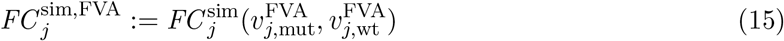

### 4.7 Software implementation

Model development was performed using Python and Jupyter notebooks with open-source Python libraries including cobrapy.^67^ The sequence of upgrades and improvements can be seen in the Git version control records. The repository is available online through Github (https://github.com/trinhlab/ctherm-gem) and in Supplementary Material 1.

### 4.8 Proteomics data collection

*C. thermocellum* wild-type and Δ*hydG*Δ*ech* strains were cultured in batch reactors and metabolic fluxes were calculated as previously described.^10^ For proteomics measurements, the wild-type and mutant strains were cultured in MNM and MTC media,^68^ respectively. While both wild-type and mutant were originally cultured in MTC,^10^ the wild-type had to be cultured separately in MNM medium due to insufficient volume for proteomics sampling in the MTC culture. MTC has higher nitrogen and trace mineral concentrations, but previous studies have shown no effect on growth rates.^68^ During the mid-exponential growth phase 10 mL samples were collected, centrifuged, and the resulting pellet was stored at −20 °C. Cell pellets were then prepared for LC–MS/MS-based proteomic analysis. Briefly, proteins extracted via SDS, boiling, and sonic disruption were precipitated with trichloroacetic acid.^69^ The precipitated protein was then resolubilized in urea and treated with dithiothreitol and iodoacetamide to reduce and block disulfide bonds prior to digestion with sequencing-grade trypsin (Sigma-Aldrich). Following two-rounds of proteolysis, tryptic peptides were salted, acidified, and filtered through a 10 kDa MWCO spin column (Vivaspin 2; GE Healthcare) and quantified by BCA assay (Pierce).

For each LC–MS/MS run, 25 μg of peptides were loaded via pressure cell onto a biphasic MudPIT column for online 2D HPLC separation and concurrent analysis via nanospray MS/MS using a LTQ-Orbitrap XL mass spectrometer (Thermo Scientific) operating in data-dependent acquisition (one full scan at 15 k resolution followed by 10 MS/MS scans in the LTQ, all one μscan; monoisotopic precursor selection; rejection of analytes with an undecipherable charge; dynamic exclusion = 30 s).^70^

Eleven salt cuts (25 mM, 30 mM, 35 mM, 40 mM, 45 mM, 50 mM, 65 mM, 80 mM, 100 mM, 175 mM and 500 mM ammonium acetate) were performed per sample run with each followed by 120 min organic gradient to separate peptides.

Resultant peptide fragmentation spectra (MS/MS) were searched against the *C. thermocellum* DSM1313 proteome database concatenated with common contaminants and reversed sequences to control false-discovery rates using MyriMatch v.2.1.^71^ Peptide spectrum matches (PSM) were filtered by IDPicker v.3^72^ to achieve a peptide-level FDR of <1 % per sample run and assigned matched-ion intensities (MIT) based on observed peptide fragment peaks. PSM MITs were summed on a per-peptide basis and those uniquely mapping to their respective proteins were imported into InfernoRDN.^73^ Peptide intensities were log_2_-transformed, normalized across replicates by LOESS, standardized by median absolute deviation, and median centered across all samples. Peptide abundance data were then assembled to proteins via RRollup and further filtered to maintain at least two values in at least one replicate set. Protein abundances were then used for the modeling efforts describe herein.

All raw and database-searched LC-MS/MS data pertaining to this study have been deposited into the MassIVE proteomic data repository and have been assigned the following accession numbers: MSV000084488 (MassIVE) and PXD015973 (ProteomeXchange). Data files are available upon publication (ftp://massive.ucsd.edu/MSV000084488/).

### 4.9 Modular cell design

The iCBI655 model in its batch reaction and cellobiose carbon source configuration (Supplementary Material 4) was used as basis for the strain design. The alcohol pathways were curated from recent literature,^6,37,74^ where adapted Adh can use either NADH or NADPH to form the target alcohol.^9^ The esters pathway require the inclusion of a condensation reaction between alcohol and acyl-CoA that are already present in the alcohol pathways. This reaction can be performed by cloramphenicol acetyl transferase (CAT) as recently demonstrated.^75^ All the software involved in the generation of these designs is available online at https://github.com/trinhlab/modcell-hpc and https://github.com/trinhlab/modcell-hpc-study.

## Acknowledgments

This research was financially supported in part by the NSF CAREER award (NSF#1553250 to CTT) and by The Center of Bioenergy Innovation (CBI), U.S. Department of Energy Bioenergy Research Center supported by the Office of Biological and Environmental Research in the DOE Office of Science (to CTT and CDM). The funders had no role in the study design, data collection and analysis, decision to publish, or preparation of the manuscript.

## Supplementary Materials

1. Software used to develop, configure, and analyze iCBI655.
2. Flux dataset used to train the iCBI655 model and proteomics dataset for the wild-type and Δ*hydG*Δ*ech* strains.
3. Supplementary tables.
4. iCBI655 model in various formats for cellobiose growth condition and map of central metabolic pathways in Escher format.

