## Supplementary Material SM1 for "Development of a genome-scale metabolic model of *Clostridium thermocellum* and its applications for integration of multi-omics datasets and strain design": README.md

Supplementary Material: 1
Study:  Development of a genome-scale metabolic model of Clostridium thermocellum and its applications for integration of multi-omics datasets and strain design
Authors: Sergio Garcia, R. Adam Thompson, Richard J. Giannone, Satyakam Dash, Costas D. Maranas, Cong Trinh.
---

This a snapshot of the Github repository containing the software used to developed, configure and analyze iCBI655. Access the online version at: https://github.com/trinhlab/ctherm-gem
