## Supplementary Material SM2 for "Development of a genome-scale metabolic model of *Clostridium thermocellum* and its applications for integration of multi-omics datasets and strain design": README.md

Supplementary Material: 2
Study:  Development of a genome-scale metabolic model of Clostridium thermocellum and its applications for integration of multi-omics datasets and strain design
Authors: Sergio Garcia, R. Adam Thompson, Richard J. Giannone, Satyakam Dash, Costas D. Maranas, Cong Trinh.
---

This S.M. contains:
- `./ctherm_extracellular_flux.csv` The flux dataset used to train the iCBI655 model
- `./07142015wt_v_hydG-ech_25pct3.xlsx` The proteomics dataset used to study the hydg-ech mutant. Additional header description:

| Number | name   | description                                 |
| ----   | ---    | --                                          |
| 1      | R1_t4  | wild type growth phase (replicate 1)        |
| 1      | R2_t5  | wild type growth phase (rep. 2 )            |
| 2      | R1_t7  | wild type stationary phase (rep. 1)         |
| 2      | R2_t9  | wild type stationary phase (rep. 2)         |
| 3      | R21_t6 | mutant (hydg-ech) growth phase (rep. 1)     |
| 3      | R22_t6 | mutant (hydg-ech) growth phase (rep. 2)     |
| 4      | R21_t9 | mutant (hydg-ech) stationary phase (rep. 1) |
| 4      | R22_t9 | mutant (hydg-ech) stationary phase (rep. 2) |

For further context and resources consult the code available in  S.M. 1 of this study and https://github.com/trinhlab/ctherm-gem
