## Supplementary Material 3 for "Development of a genome-scale metabolic model of *Clostridium thermocellum* and its applications for integration of multi-omics datasets and strain design"

Table S1: The 70 consistent reactions in the $\Delta hydG \Delta ech$ case study and their associated fold changes. The biomass reaction is not included due to size. Note that this table is continued in the following pages.

| ID | Formula | Fold change | | |
|---|---|---|---|---|
| | | *proteomics* | *pFBA* | *FVAcenter* |
| MDH | mal__L_c + nad_c ↔ h_c + nadh_c + oaa_c | 1.4 | 0.2 | 0.1 |
| PEPCK_re | co2_c + gdp_c + pep_c → gtp_c + oaa_c | 0.9 | 0.1 | 0.1 |
| VOR2b | 3mob_c + coa_c + 2.0 fdxo_42_c → co2_c + 2.0 fdxr_42_c + h_c + ibcoa_c | 0.9 | 12.3 | 0.8 |
| PFL | coa_c + pyr_c → accoa_c + for_c | 0.5 | 0.2 | 0.8 |
| FRNDPR2r | 2.0 fdxr_42_c + h_c + nadh_c + 2.0 nadp_c ↔ 2.0 fdxo_42_c + nad_c + 2.0 nadph_c | 0.3 | 0.2 | 2.3 |
| ALCD2x | acald_c + h_c + nadh_c → etoh_c + nad_c | 0.1 | 0.8 | 0.6 |
| IBUTCOARx | h_c + ibcoa_c + nadh_c → 2mppal_c + coa_c + nad_c | 0.1 | 12.3 | 0.8 |
| ACALD | accoa_c + h_c + nadh_c → acald_c + coa_c + nad_c | 0.1 | 0.8 | 1.1 |
| ALCD23xi | 2mppal_c + h_c + nadh_c → ibutoh_c + nad_c | 0.1 | 12.3 | 0.8 |
| ME2 | mal__L_c + nadp_c → co2_c + nadph_c + pyr_c | 0.1 | 0.3 | 0.1 |
| PSCVT | pep_c + skm5p_c → 3psme_c + pi_c | -0.1 | -0.3 | 0.7 |
| PTAr | accoa_c + pi_c ↔ actp_c + coa_c | -0.1 | -2.7 | -0.1 |
| HSOR | 3.0 h_c + 3.0 nadph_c + so3_c → 3.0 h2o_c + h2s_c + 3.0 nadp_c | -0.1 | -0.3 | -1.2 |
| TRDR | h_c + nadph_c + trdox_c → nadp_c + trdrd_c | -0.1 | -0.3 | 0.8 |
| IGPS | 2cpr5p_c + h_c → 3ig3p_c + co2_c + h2o_c | -0.1 | -0.3 | 0.2 |
| GLUDy | glu__L_c + h2o_c + nadp_c ↔ akg_c + h_c + nadph_c + nh4_c | -0.1 | -0.5 | -0.2 |
| ECH | 2.0 fdxr_42_c + 3.0 h_c ↔ 2.0 fdxo_42_c + h2_c + h_e | -0.1 | -15.6 | -13.5 |
| ALLAS | 24.0 ala__D_c + 24.0 atp_c + 24.0 cdpglyc_c + dg12dg_c + 24.0 h2o_c → ala_lta_c + 24.0 amp_c + 24.0 cmp_c + 48.0 h_c + 24.0 ppi_c | -0.1 | -0.2 | 0.0 |
| HSDxi | aspsa_c + h_c + nadh_c → hom__L_c + nad_c | -0.1 | -0.3 | 0.8 |
| PRMICI | prfp_c → prlp_c | -0.1 | -0.3 | -0.1 |
| OCBT | cbp_c + orn__L_c ↔ citr__L_c + h_c + pi_c | -0.2 | -0.3 | -0.3 |
| UAGCVT | pep_c + uacgam_c → pi_c + uaccg_c | -0.2 | -0.3 | -0.1 |
| ANS | chor_c + gln__L_c → anth_c + glu__L_c + h_c + pyr_c | -0.2 | -0.3 | 0.2 |
| DHAD2 | 23dhmp_c → 3mop_c + h2o_c | -0.2 | -0.3 | 0.6 |
| ASPO2y | asp__L_c + nadp_c → h_c + iasp_c + nadph_c | -0.3 | -0.3 | -0.1 |
| NADK | atp_c + nad_c → adp_c + h_c + nadp_c | -0.3 | -0.3 | -0.1 |

| ID | Formula | Fold change | | |
|---|---|---|---|---|
| | | *proteomics* | *pFBA* | *FVAcenter* |
| METS | 5mthf_c + hcys__L_c → met__L_c + thf_c | -0.3 | -0.3 | 0.8 |
| IMPD | h2o_c + imp_c + nad_c ↔ h_c + nadh_c + xmp_c | -0.3 | -0.3 | -0.3 |
| MG2abc | atp_c + h2o_c + mg2_e → adp_c + h_c + mg2_c + pi_c | -0.3 | -0.3 | -0.1 |
| PDHam1hi | h_c + pyr_c + thmpp_c → 2ahethmpp_c + co2_c | -0.4 | -0.3 | 12.4 |
| ACAS_2ahbut | 2ahethmpp_c + 2obut_c → 2ahbut_c + thmpp_c | -0.4 | -0.3 | 0.6 |
| GF6PTA | f6p_B_c + gln__L_c → gam6p_c + glu__L_c | -0.4 | -0.3 | 0.0 |
| CHRS | 3psme_c → chor_c + pi_c | -0.4 | -0.3 | 0.7 |
| SERH | 3ig3p_c + ser__L_c → g3p_c + h2o_c + trp__L_c | -0.4 | -0.3 | 0.2 |
| ALATA_L | akg_c + ala__L_c ↔ glu__L_c + pyr_c | -0.5 | -0.3 | -12.9 |
| NNDPR | h_c + prpp_c + quln_c → co2_c + nicrnt_c + ppi_c | -0.5 | -0.3 | -0.1 |
| TMDS | dump_c + mlthf_c → dhf_c + dtmp_c | -0.5 | -0.3 | -0.1 |
| PHETA1 | akg_c + phe__L_c ↔ glu__L_c + phpyr_c | -0.6 | -0.3 | 0.8 |
| TYRTA | akg_c + tyr__L_c ↔ 34hpp_c + glu__L_c | -0.6 | -0.3 | 0.6 |
| GMPS | atp_c + nh4_c + xmp_c → amp_c + gmp_c + 3.0 h_c + ppi_c | -0.6 | -0.3 | -0.3 |
| SKK | atp_c + skm_c → adp_c + h_c + skm5p_c | -0.6 | -0.3 | 0.7 |
| ACOTA | acorn_c + akg_c ↔ acg5sa_c + glu__L_c | -0.6 | -0.3 | -0.3 |
| PPDK | amp_c + 2.0 h_c + pep_c + ppi_c → atp_c + pi_c + pyr_c | -0.7 | -10.0 | 0.1 |
| GLUPRT | gln__L_c + h2o_c + prpp_c → glu__L_c + h_c + ppi_c + pram_c | -0.7 | -0.3 | -0.3 |
| KARI | 2ahbut_c ↔ cpd10162_c | -0.8 | -0.3 | 0.6 |
| KARI_23dhmp | 23dhmp_c + nadp_c ↔ cpd10162_c + h_c + nadph_c | -0.8 | -0.3 | 0.6 |
| ARGSL | argsuc_c ↔ arg__L_c + fum_c | -0.8 | -0.3 | -0.3 |
| LEUTA | 4mop_c + glu__L_c → akg_c + leu__L_c | -0.8 | -0.3 | 0.4 |
| ILETA | akg_c + ile__L_c ↔ 3mop_c + glu__L_c | -0.8 | -0.3 | 0.6 |
| VALTA | akg_c + val__L_c ↔ 3mob_c + glu__L_c | -0.8 | -1.4 | -1.5 |
| ARGSS | asp__L_c + atp_c + citr__L_c ↔ amp_c + argsuc_c + 2.0 h_c + ppi_c | -0.8 | -0.3 | -0.3 |
| SHSL2 | h2s_c + suchms_c → hcys__L_c + succ_c | -0.9 | -6.0 | 0.8 |
| AHSL | achms_c + cys__L_c ↔ ac_c + cyst_L_c + h_c | -0.9 | -10.6 | -0.3 |
| SHSL1 | cyst_L_c + h_c + succ_c ↔ cys__L_c + suchms_c | -0.9 | -10.6 | 0.4 |
| ACKr | actp_c + adp_c → ac_c + atp_c | -0.9 | -2.7 | -0.1 |
| QULNS | dhap_c + iasp_c → 2.0 h2o_c + h_c + pi_c + quln_c | -0.9 | -0.3 | -0.1 |

| ID | Formula | Fold change | | |
|---|---|---|---|---|
| | | *proteomics* | *pFBA* | *FVAcenter* |
| NADS2 | atp_c + dnad_c + gln__L_c + h2o_c → amp_c + glu__L_c + 2.0 h_c + nad_c + ppi_c | -0.9 | -0.3 | -0.1 |
| FE3abc | atp_c + fe3_e + h2o_c → adp_c + fe3_c + h_c + pi_c | -1.0 | -0.3 | -0.1 |
| ASPTA | akg_c + asp__L_c ↔ glu__L_c + oaa_c | -1.0 | -0.3 | 0.2 |
| CTPS1 | atp_c + nh4_c + utp_c → adp_c + ctp_c + 2.0 h_c + pi_c | -1.2 | -0.3 | 0.0 |
| ACGK | acglu_c + atp_c → acg5p_c + adp_c | -1.2 | -0.3 | -0.3 |
| IGPDH | eig3p_c → h2o_c + imacp_c | -1.2 | -0.3 | -0.1 |
| AGPR | acg5sa_c + nadp_c + pi_c ↔ acg5p_c + h_c + nadph_c | -1.3 | -0.3 | -0.3 |
| PHEt2r | h_e + phe__L_e ↔ h_c + phe__L_c | -1.5 | -0.3 | 0.8 |
| UAG4Ei | uacgam_c → udpacgal_c | -1.5 | -0.3 | -0.1 |
| CYSS | acser_c + h2s_c → ac_c + cys__L_c | -1.8 | -0.3 | -0.5 |
| BIF | 2.0 fdxr_42_c + 3.0 h_c + nadh_c ↔ 2.0 fdxo_42_c + 2.0 h2_c + nad_c | -1.8 | -13.8 | -12.5 |
| UMPK | atp_c + h_c + ump_c → adp_c + udp_c | -2.1 | -0.3 | 0.0 |
| SULabc | atp_c + h2o_c + so4_e → adp_c + h_c + pi_c + so4_c | -4.6 | -0.3 | 0.8 |

Table S2: The pFBA-simulated fluxes from the $\Delta hydG \Delta ech$ case study. For this table, we only presented reactions involving NADPH or exchange reactions that have different fluxes between wild-type and mutant. We did not present the biomass reaction due to size as well as the reactions with the same fold change magnitude as the biomass reaction ($|FC| = 0.26$), likely because they are fully correlated.

| ID | Formula | Fluxes (mmol/gCDW/hr) | | |
|---|---|---|---|---|
| | | *W.T.* | *Mut.* | FC |
| EX_ibutoh_e | ibutoh_e → | 0.0 | 0.49 | 12.26 |
| KARA1 | alac__S_c + h_c + nadph_c → 23dhmb_c + nadp_c | 0.22 | 0.59 | 1.42 |
| EX_etoh_e | etoh_e → | 1.09 | 1.88 | 0.78 |
| EX_h2o_e | h2o_e ↔ | -0.38 | 0.5 | 0.42 |
| EX_co2_e | co2_e → | 2.07 | 2.77 | 0.42 |
| ME2 | mal__L_c + nadp_c → co2_c + nadph_c + pyr_c | 2.9 | 3.51 | 0.28 |
| EX_for_e | for_e → | 0.48 | 0.56 | 0.23 |
| FRNDPR2r | 2.0 fdxr_42_c + h_c + nadh_c + 2.0 nadp_c ↔ 2.0 fdxo_42_c + nad_c + 2.0 nadph_c | -1.02 | -1.17 | 0.2 |
| EX_nh4_e | nh4_e ↔ | -0.73 | -0.53 | -0.48 |
| GLUDy | glu__L_c + h2o_c + nadp_c ↔ akg_c + h_c + nadph_c + nh4_c | -0.61 | -0.42 | -0.53 |
| EX_h_e | h_e ↔ | 2.93 | 1.19 | -1.3 |
| EX_val__L_e | val__L_e → | 0.18 | 0.06 | -1.54 |
| ICDHyr | icit_c + nadp_c → akg_c + co2_c + nadph_c | 0.21 | 0.04 | -2.27 |
| EX_ac_e | ac_e → | 0.93 | 0.17 | -2.46 |
| EX_lac__L_e | lac__L_e → | 0.05 | 0.01 | -2.49 |
| EX_succ_e | succ_e → | 0.41 | 0.0 | -12.01 |
| EX_h2_e | h2_e → | 2.2 | 0.0 | -14.43 |
