## Supplementary Material 4 for "Development of a genome-scale metabolic model of *Clostridium thermocellum* and its applications for integration of multi-omics datasets and strain design": README.md

Supplementary Material: 4
Study: Development of an updated genome-scale metabolic model of \textit{Clostridium thermocellum} and its application for integration of multi-omics datasets
Authors: Sergio Garcia, R. Adam Thompson, Richard J. Giannone, Satyakam Dash, Costas D. Maranas, Cong Trinh.
---
This S.M. contains:
-  The iCBI655 model in various formats for cellobiose growth condition. For additional context and tools to change model settings refer to the code*.
- The Escher metabolic map (https://escher.github.io) of central metabolism to visualize model results. For additional maps refer to the code*.

* The code is available in  S.M. 1 of this study and https://github.com/trinhlab/ctherm-gem
